# Strengthening Bolivian Pharmacovigilance system: new therapeutic strategies to improve health of Chagas Disease and Tuberculosis patients

**DOI:** 10.1101/441949

**Authors:** Nuria Cortes-Serra, Ruth Saravia, Rosse Mary Grágeda, Amílcar Apaza, Jorge Armando González, Brenda Ríos, Joaquim Gascon, Faustino Torrico, Maria-Jesus Pinazo

**Author notes:** Correspondence to: **Nuria Cortes-Serra.** Instituto de Salud Global de Barcelona (ISGlobal), Hospital Clínic i Provincial/ IDIBAPS, Universitat de Barcelona.

## Abstract

**Introduction:** Chagas disease (CD) and Tuberculosis (TB) are important health problems in Bolivia. Current treatments for both infections require a long period of time, and unwanted drug-related adverse events (ADRs) are frequent.

**Purpose:** This study aims to strengthen the Bolivian Pharmacovigilance system, focusing on CD and TB.

**Methods:** A situational diagnosis of Pharmacovigilance in the Department of Cochabamba was performed. The use of a new Local Case Report Form (CRF) was implemented, together with the CRF established by the Unidad de Medicamentos y Tecnología en Salud (UNIMED), in several health care centers. Training and follow-up on drug safety monitoring and ADR reporting was provided to all health professionals involved in CD and TB treatment. A comparative analysis of the reported ADRs using the CRF provided by UNIMED, the new CRF proposal, and medical records, was performed.

**Results:** Out of the total patients starting treatment for CD, 35,35% suffered ADR according to the information collected in the medical records, and 25% of them were classified as moderate/severe (MS) types. Only 51,43% of MS ADRs were reported to UNIMED. Regarding TB treatment, 9,89% of the total patients suffered ADR, 44% of them were classified as MS, and 75% of MS ADRs were reported to UNIMED.

**Conclusions:** The reinforcement of the Bolivian Pharmacovigilance system is an ambitious project that should take a long-term perspective and the engagement of national health workers and other stake holders at all levels. Continuity and perseverance are essential to achieve a solid ADR reporting system, improving patient safety, drug efficacy and adherence to treatment.

## 1. INTRODUCTION

Chagas disease (CD), caused by the parasite *Trypanosoma cruzi (T. cruzi)*, is one of the main health problems in Latin America. *T. cruzi* infection has been declared as a major public health issue in the region, affecting approximately 6 to 7 million people worldwide. Bolivia is the country with the highest prevalence and incidence of CD. In addition, travel and immigration patterns have increased the relevance of *T. cruzi* infection outside of endemic areas [1–3].

CD is recognized as one of the 17 neglected tropical diseases (NTDs), for which pharmaceutical industry does not assign resources to invest in research and development of new drugs. Approved drugs for treatment of *T. cruzi* infection (beznidazole and nifurtimox) are complex, and unwanted drug-related adverse events (ADRs) are frequent. The onset of the ADR is one of the main causes that lead to patient abandonment, resulting in therapeutic failure or ineffective treatment [4–9].

Tuberculosis (TB) is another important health problem in Bolivia, where the prevalence and incidence of the disease is high compared to other South American countries. TB treatment involves the combination of several drugs over a long period of time, with high frequency of ADRs, which compromise the effectiveness of treatment. In addition, resistance to current drugs has been increasing during the last years [10–12].

In the management of CD and TB, poor adherence to treatment continues to be another challenge to face. Factors contributing to poor medication adherence include lack of follow-up protocols during treatment, and toxicity of the drugs. Nevertheless, with an adequate management of the drugs, most of patients are able to finish treatment even in case of ADRs, which in CD are usually mild and well controlled with symptomatic treatment. Close medical follow-up, monitoring ADRs and implementation of robust Pharmacovigilance systems are essential factors to avoid patient abandonment and achieve therapeutic success [13–17].

In the current context of CD and TB in Bolivia, strengthening the Bolivian Pharmacovigilance system is proposed as a key issue to reinforce therapeutic strategies. To implement a strong and consolidated Pharmacovigilance system including local health centers will allow to better detect and report ADRs in order to provide knowledge to improve patient safety and adherence of treatments.

## 2. METHODOLOGY OF INTERVENTION

### 2.1 Actors involved

The current project has been implemented by Barcelona Institute for Global Health (ISGlobal) and Fundación Ciencia y Estudios Aplicados para el Desarrollo en Salud y Medio Ambiente (CEADES). Both institutions shape the Bolivian Chagas Platform, a model to protocolize together with the Chagas National and Departmental Program the attention for adults with *T. cruzi* infection, and, by this way, to increase the number of adult patients diagnosed and treated for CD [17]. The research was done in collaboration with the Pharmacovigilance Unit (UNIMED) of the Bolivian Ministry of Health, the Pan American Health Organization (PAHO), the Departmental Chagas Program of Cochabamba (ChDP) and the Tuberculosis Departmental Program of Cochabamba (TBDP). A total of fourteen rural and urban primary and secondary health care centers participated in the study, including four health care centers of the Bolivian Chagas Platform.

### 2.2 Intervention plan

The intervention plan was performed in several steps:

#### A - Situational diagnosis

A situational diagnosis of Pharmacovigilance in the Department of Cochabamba was performed in order to deeply study the causes of lack of reporting ADRs. The data collection method was the use of a survey which included questions related with knowledge and attitude towards Pharmacovigilance. All health professionals involved in CD treatment with at least one year of experience working in the national health system were selected to participate. The decision to include TB in the project was proposed by UNIMED after performing the situational diagnosis.

#### B - Strategies to reinforce the current Bolivian Pharmacovigilance System

Strengthening the Bolivian Pharmacovigilance system was proposed in our project as a key issue to reinforce therapeutic strategies in CD and TB. The Bolivian Pharmacovigilance system is based on fill out the CRF elaborated by UNIMED by health care professionals. Then, the form is sent to the healthcare network management, and finally it arrives to UNIMED. Together with local health entities and UNIMED, the implementation of a new Local Case Report Form (CRF) was suggested to the Bolivian Ministry of Health. The new form was elaborated based on the follow-up form used in the Bolivian Chagas Platforms, the CRF established in the national health system by UNIMED, and other CRFs used in other countries.

The CRF elaborated by UNIMED and the new CRF proposal were implemented in the Bolivian Chagas Platforms located in the Department of Cochabamba (one in urban area and three in rural areas), and in the primary and secondary national health care centers conforming the network. The CRF provided by UNIMED was already established in some of the health care facilities. Both forms were available in physical and electronic format, which was assigned randomly in each health care center. To evaluate the intervention, a comparative analysis of the use of the CRF established by UNIMED, the new CRF proposal, and the medical records to report ADRs was performed. The medical records of the total patients starting treatment for CD or TB were analyzed in order to detect how many of them suffered ADR during the treatment.

A specific training on drug safety monitoring and ADR reporting was given, together with PAHO and UNIMED, to all health professionals involved in CD and TB treatment. Three follow-up visits were performed in each health care center.

#### C - Analysis of the ADRs reported

An analysis of the ADRs reported was performed, focusing on the following variables: total patients reporting ADR; total patients abandoning treatment; affected organ-system; severity of ADRs reported; recurrents ADRs; health care intervention; differences in reporting rates between the CRF established by UNIMED, the new CRF proposal and the ADRS reported in the medical records; differences in reporting rates between physical and electronic CRF format; and differences in reporting rates between health care centers.

Patient abandonment was defined as the failure to complete medically indicated curative therapy.

#### D - Data analysis

Absolute and relative frequency counts, and measures of central tendency (mean) were calculated. GraphPad PRISM 5.0 software was used to draw graphs.

#### E - Ethics statement

The study protocol was approved by the Ethics Committee of the Hospital Clinic of Barcelona and the Ethics Committee of CEADES.

## 3. RESULTS

### 3.1 Situational diagnosis: a challenging beginning

Health care professionals confirmed the importance of reporting ADRs, giving an average score of 9,6 out of 10. This figure contrasts with the effectively reported ADR rate: 4,74 out of 10.

Causes for underreporting ADRs by health professionals were multifaceted. The most common reason for not reporting ADRs was “lack of knowledge and awareness about Pharmacovigilance and the duty of reporting” (50%). Health professionals also reported “lack of availability of report forms” (18,75%), “forgetfulness to fill out the form” (12,5%), “forms with too many variables” (6,25%), “patients forgetting to mention the ADR” (6,25%) and “ADRs with no clinical significance” (6,25%). Results are summarized in Fig 1.

**Figure 1:**
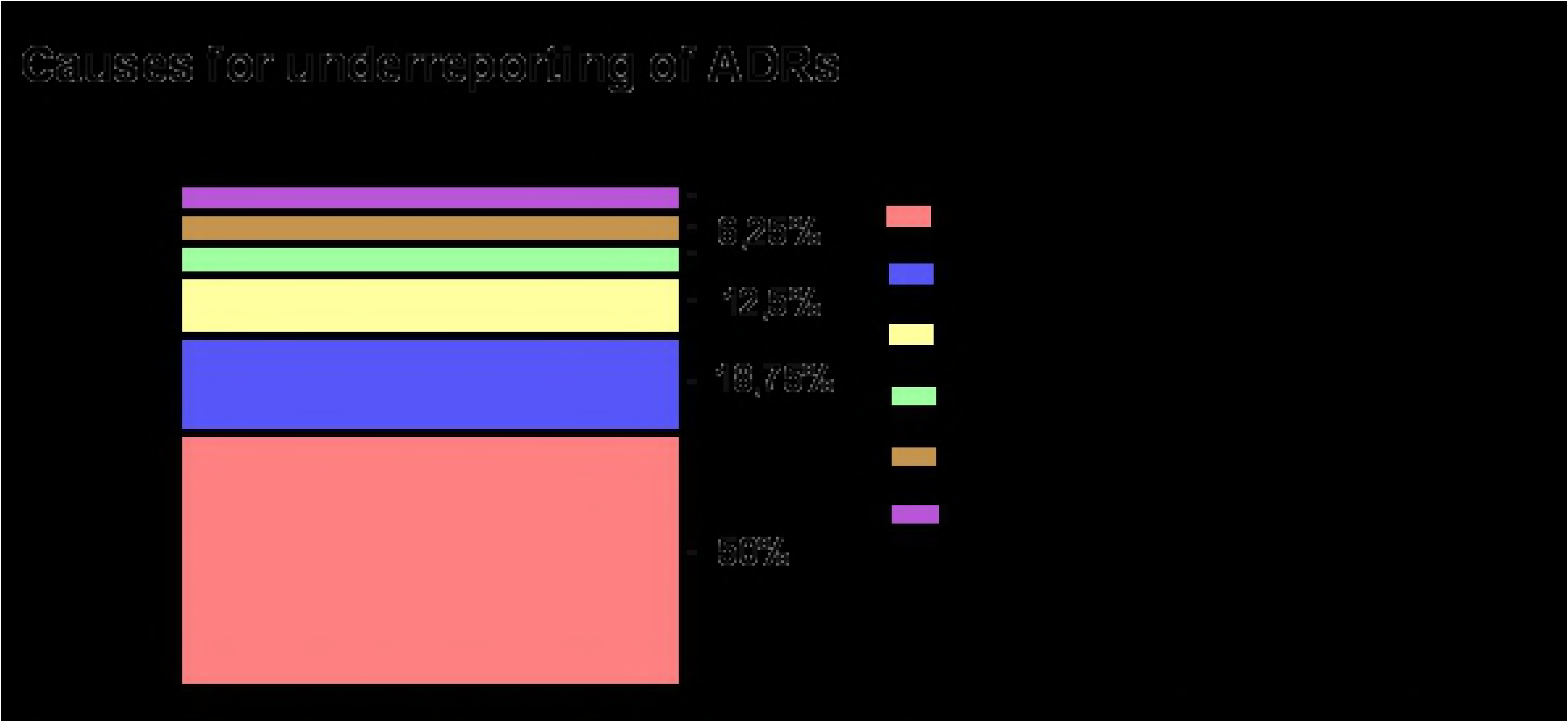
Causes for underreporting of ADRs.

From all the health care professionals participating in the survey, 53% reported ADRs using patient’s medical record, 27% using CRFs established by UNIMED, and 20% using CRFs obtained from Bolivian ChDP. Regarding the CRF established by UNIMED, 62,5 % of health care professionals were not aware of it. Of the 37,5% that were aware about the CRF from UNIMED, only 33,3% of them were able to fill out the form correctly.

In terms of health care workers opinion, the most appropriate method to report ADRs was the use of electronic CRFs (50%), the use of CRFs in physical format (31,25%) or irrelevant (18,75%).

Regarding the issue of which ADRs should be reported to UNIMED, 43,75% of the health care professionals participating in our study considered important to report all type of ADRs, 31,25% to report moderate and severe ADRs, and 25% to report only severe ADRs.

Lack of feedback from UNIMED to health care professionals could be another cause for underreporting of ADRs. A huge percentage of health care professionals (81,25%) considered that receiving feedback from UNIMED would increase the reporting rate of ADRs.

### 3.2 Evaluation of the intervention

From the 396 patients starting treatment for CD, 140 (35,35% of them) suffered ADR according to the information collected in the medical records. Out of the total ADRs collected in the medical records, 25% of them were classified as moderate or severe types, and should have been reported to the Bolivian Pharmacovigilance system. Only 51,43% of them were reported to UNIMED.

Regarding TB treatment, 9,89% of the total patients suffered ADR. Out of the total ADRs collected in the medical records, 44% of them were classified as moderate or severe types, and
should have been reported to the Bolivian Pharmacovigilance system. From all the ADRs that should have been reported, 75% of them were reported to UNIMED.

Results are summarized in Fig 2.

**Figure 2:**
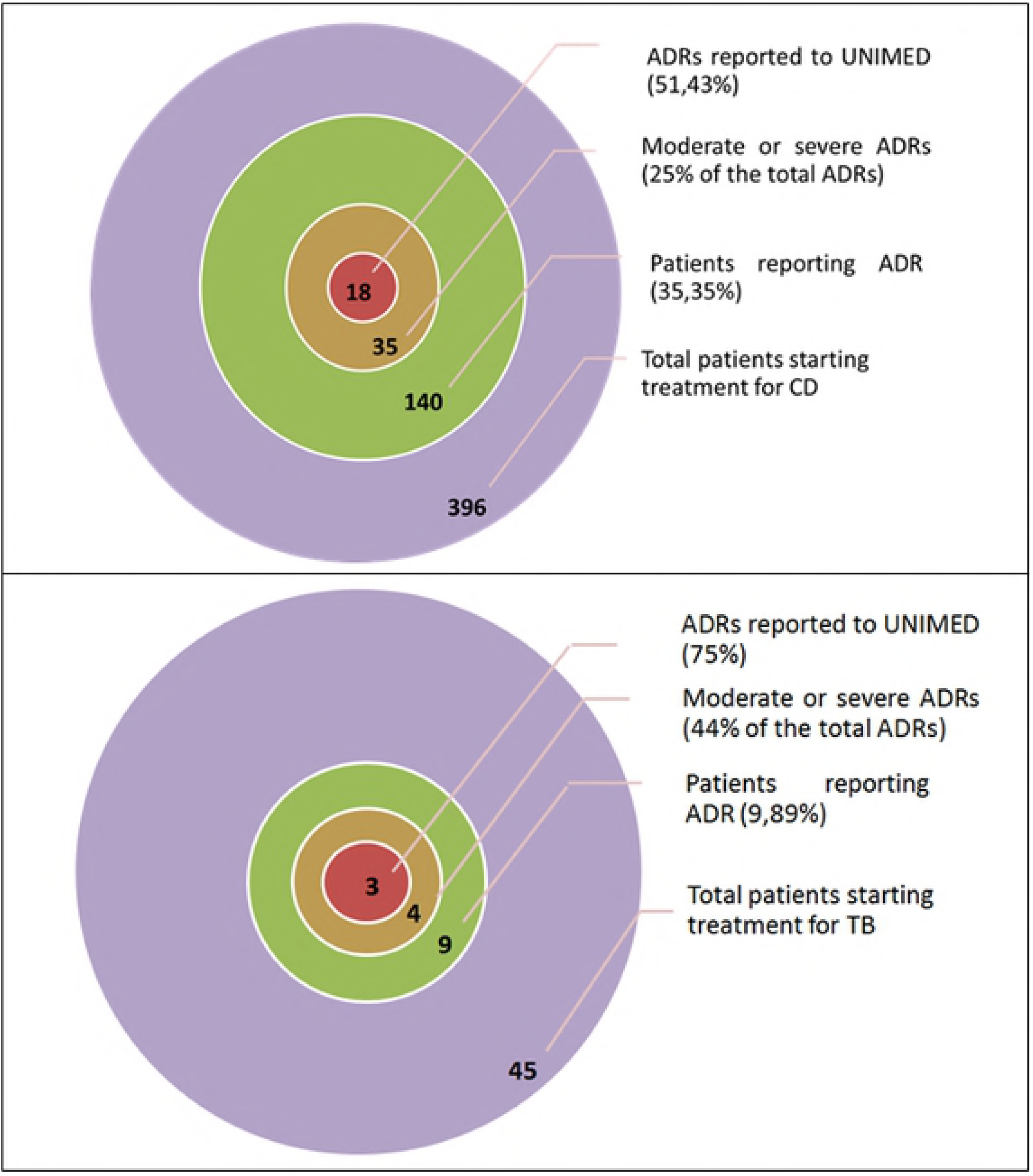
Summary of the intervention results.

### 3.3 Analysis of the ADRs reported

From a total of 396 patients participating in the study and starting treatment for CD, 140 (35,35%) reported with ADRs. Patient abandonment was of 12,63%. Results are shown in table 1.

**Table 1:** Total patients starting treatment, reporting ADRs, and abandoning treatment for CD in the Bolivian Chagas Platforms located in the Department of Cochabamba and the primary and secondary national health care centers included in the study.

|  | Cercado network | Sacaba network | Punata network | Villatunari network | Total |
| --- | --- | --- | --- | --- | --- |
| Total patients starting treatment | 198 | 47 | 59 | 92 | 396 |
| Total patients reporting ADR | 89 (50%) | 22 (46,81%) | 10 (16,95%) | 19 (20,65%) | 140 (35,35%) |
| Total patients abandoning treatment | 24 (15,66%) | 10 (21,28%) | 5 (8,47%) | 11 (11,96%) | 50 (12,63%) |

From 91 patients starting treatment for TB, 9 (9,89%) reported with ADRs, and the percentage of patient abandonment was 2,20%. Results are shown in table 2.

Concerning CD, 35% of the total ADRs presented during treatment were dermatological, 29% affected the central nervous system, 20% were gastrointestinal, and 16% affected other organs or systems. Most of the ADRs presented during CD treatment were mild (74%), whereas 15% were classified as mild/moderate, 10% as moderate, and 1% as severe. When focusing on the health care intervention, nearly all ADRs were treated in the primary health care center (99%), and only 1% had to be referred. Finally, the majority of the ADRs were non-recurrent (72%), while 28% were recurrent ADRs.

**Table 2:**
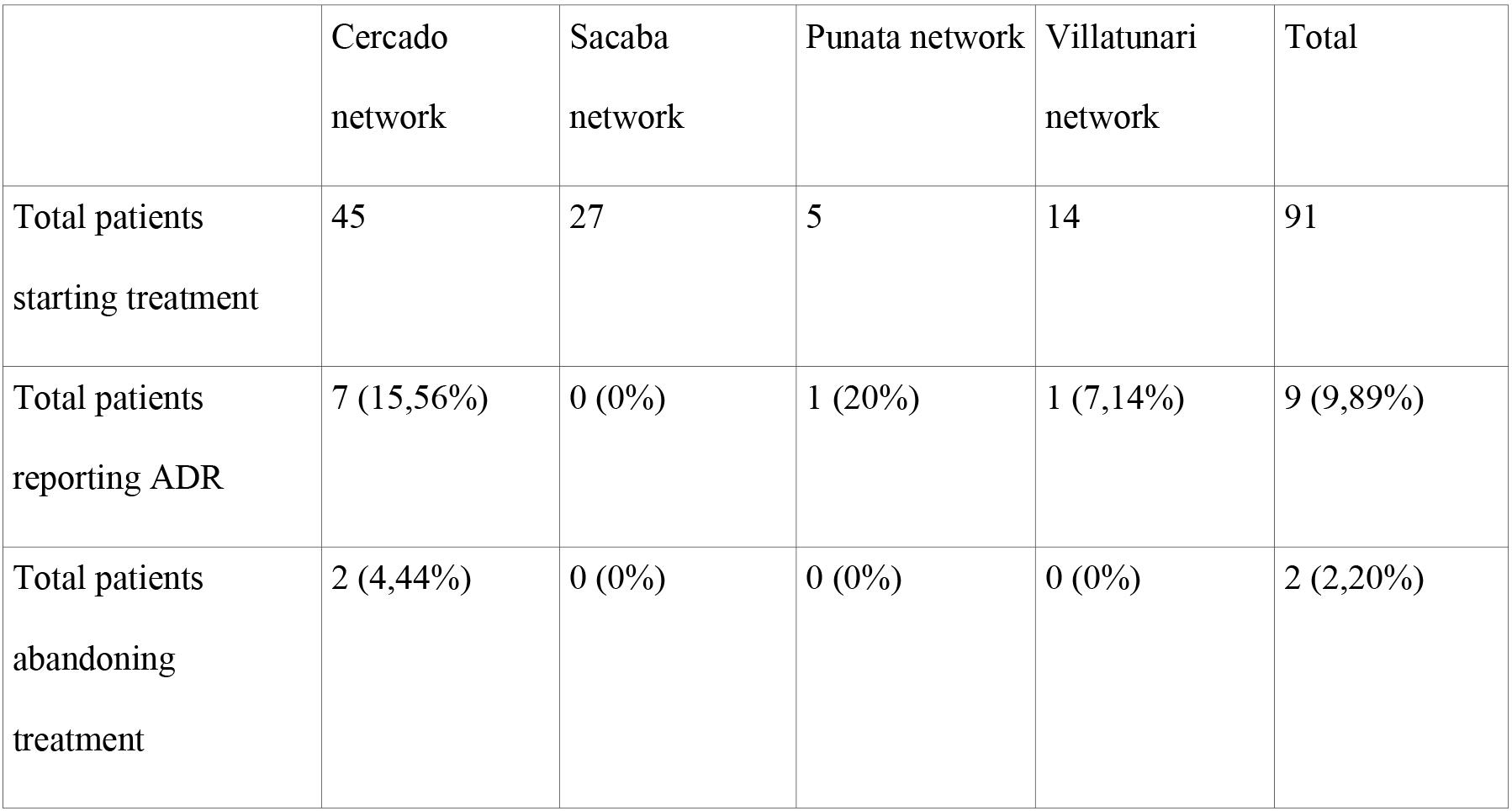
Total patients starting treatment, reporting ADRs, and abandoning treatment for TB in the primary and secondary national health care centers included in the study.

|  | Cercado network | Sacaba network | Punata network | Villatunari network | Total |
| --- | --- | --- | --- | --- | --- |
| Total patients starting treatment | 45 | 27 | 5 | 14 | 91 |
| Total patients reporting ADR | 7 (15,56%) | 0 (0%) | 1 (20%) | 1 (7,14%) | 9 (9,89%) |
| Total patients abandoning treatment | 2 (4,44%) | 0 (0%) | 0 (0%) | 0 (0%) | 2 (2,20%) |

Regarding the characteristics of the ADRs presented during TB treatment, 36,36% of the total ADRs presented were dermatological, 36,36% gastrointestinal, and 27,27% affected the central nervous system. In terms of severity, 56% of the ADRs presented were classified as mild, 33% were considered moderate, and 11% were severe. Regarding health care intervention, 56% of the ADRs presented were treated in the primary health care center, 33% were referred, and 11% were not reported. Lastly, most of the ADRs presented were non-recurrent (78%), while 22% of them were recurrent.

## 4. DISCUSSION

Pharmacovigilance programme is an important component of National Healthcare Systems (NHS). Data coming from Pharmacovigilance programme is essential to ensure safety and effectiveness of drugs and to provide information concerning regulatory actions [18]. The efforts of the Bolivian Ministry of Health to increase access to treatment for CD and TB require strengthening the Pharmacovigilance system.

Despite having developed tools to report ADRs, Bolivian Pharmacovigilance System was not well known for most health professionals up to now. Together with UNIMED, the ChDP and the TBDP, an extended lack of knowledge and awareness about Pharmacovigilance was detected, as well as the duty of reporting, and the lack of reporting ADR forms provided by UNIMED. Insufficient training for healthcare professionals was the most important and urgent weakness detected, especially concerning the primary national health care centers located in rural areas. Training and follow-up on drug safety monitoring and ADR reporting was performed to all health professionals involved in CD and TB treatment. A more specific training on Pharmacovigilance was provided to all health professionals working in the national health care centers of rural areas (departments of Villatunari and Punata).

Although our project has shown interesting results, it is important to highlight that to reinforce the Bolivian Pharmacovigilance System requires persistence and an active follow-up with actors directly implicated in the NHS. This study emphasizes the importance of an educational intervention to change attitude towards ADR reporting among healthcare professionals. However, since it was a particular intervention, its long-term effect could not be measured. Out of the total moderate and severe ADRs presented during CD treatment, only half of them were reported to the Bolivian Pharmacovigilance system. Regarding TB, 25% of the moderate and severe ADRs presented were not reported to the Bolivian Pharmacovigilance system. It is important to notice that health care professionals are more used to report ADRs related to TB than to CD, even though CD has been declared as a health priority for the Ministry of Health [19]. TBDP had a pioneering role in promoting national ADR reporting system and providing training for health care professionals [20]. Due to the characteristics of the current therapeutic schemes and the scaling-up of Chagas treatment, ChDP should put more emphasis on Pharmacovigilance activities to help physicians in their clinical practice.

The percentage of patient abandonment was 12,63% in CD patients, and 2,20% in TB patients. It is highlighted that although abandonment levels may seem not to be high in TB, TB treatment abandonment has fatal consequences for patients, which also become potential sources of infection and resistance to available drugs [21].

Differences in reporting rates between the CRF established by UNIMED and the new CRF proposal were not found. However, the new proposal of CRF results in better quality of data collected. Information about the characteristics of presented ADRs in terms of severity, affected organ-system, clinical suspicion of recurrence, and health care intervention is collected in the CRF proposal. Better designed, user-friendly and standardized reporting forms would improve the process of capturing accurate information about ADR events [18]. New tools adapted to the reality of health care workers are needed in order to strengthen the current Bolivian ADR reporting system.

Lack of internet access is still an important problem in some regions of Bolivia, especially in rural areas. In several rural and urban health care centers included in our study internet connectivity was limited, and it was not possible to implement the CRF in electronic format. Nevertheless, other possibilities could be explored in order to overcome this limitation. Apps or electronic devices working in and offline mode could be an alternative [22]. Besides, some of the health care facilities participating in the study are currently under renovation, and it was considered important to have both forms (the CRF established by UNIMED and the new CRF proposal) available in electronic format in order to facilitate reporting.

Lack of knowledge by health care professionals regarding the classification of ADRs by severity was detected during the study. In several medical records, it was found the category «mild / moderate», which is not correct. Lack of consensus on definition of ADR was also detected, which is important also for the clinical management of the ADRs. In several medical records, symptoms experimented by patients probably associated with concomitants pathologies presented during CD or TB treatment were considered as ADRs. During our study, it was considered as ADR every event classified as that in medical records. Nevertheless, there is a need of overcoming these limitations.

Even if strengthen Bolivian Pharmacovigilance System has proven to be a good strategy to improve patient health and increase adherence to CD and TB treatment, currently the system still presents some weaknesses. According to all actors involved in our project, the implementation of the next policies was suggested:

- To strengthen Pharmacovigilance activities in UNIMED, with external funding sources if it is required, at least in a preliminary phase.
- To provide continued training in Pharmacovigilance for health care professionals, validated by the Bolivian Ministry of Health and the Pan American Health Organization.
- To perform a continuous surveillance in Pharmacovigilance activities in health care centers through monitoring and evaluation programs provided by UNIMED. In a preliminary stage, this could require external technical support or funding sources.
- To integrate the topic Pharmacovigilance into the curricula in medical, pharmacy and nursing schools, to provide excellent training to future health care professionals.

## 5. CONCLUSIONS

Overall, our findings suggest that Bolivian Pharmacovigilance system still presents some challenges that should be addressed in the next years in order to achieve a strong, integrated and consolidated ADR reporting system. The reinforcement of the Bolivian Pharmacovigilance system is an ambitious project that should take a long-term perspective and several steps. Continuity and perseverance are essential to achieve a solid and integrated reporting of ADR system. UNIMED’s current approach is to strengthen Bolivian Pharmacovigilance System in all care levels, focusing on other neglected and prevalent pathologies in Bolivia and including all the departments of the country. Several of our proposals are already in the process of being implemented. However, the responsibility of leading these actions is still unclear. Further projects are needed in order to achieve a strong and consolidated ADR reporting system to improve patient safety, drug efficacy and adherence to treatment. A medium and long-term follow up to evaluate their impact is also required.

## 6. ACKNOWLEDGEMENTS

We thank Leire Pajín for her support, critical review and perspective, and all bolivian health personnel implied in the project.

